# Intermittent hypoxia therapy engages multiple longevity pathways to double lifespan in *C.elegans*

**DOI:** 10.1101/2022.10.13.512140

**Authors:** Jason N. Pitt, Eduardo Chavez, Kathryn M. Provencher, Michelle Chen, Christina Tran, Jennifer Tran, Karen Huang, Anuj Vaid, Marian L Abadir, Naheed Arang, Scott F. Leiser, Mark B. Roth, Matt Kaeberlein

## Abstract

Genetic activation of the hypoxia response robustly extends lifespan in *C. elegans*, while environmental hypoxia shows more limited benefit. Here we describe an intermittent hypoxia therapy (IHT) able to double the lifespan of wildtype worms. The lifespan extension observed in IHT does not require HIF-1 but is partially blocked by loss of DAF-16/FOXO. RNAseq analysis shows that IHT triggers a transcriptional state distinct from continuous hypoxia and affects down-stream genes of multiple longevity pathways. We performed a temperature sensitive forward genetic screen to isolate mutants with delayed nuclear localization of DAF-16 in response to IHT and suppression of IHT longevity. One of these mutations mapped to the enzyme Inositol Polyphosphate MultiKinase (IPMK-1). *ipmk-1* mutants, like *daf-16* mutants, partially suppress the benefits of IHT, while other effectors of phosphatidyl inositol signaling pathways (PLCβ4, IPPK, Go/iα) more robustly suppress IHT longevity.

**One-Sentence Summary:** Intermittent hypoxia therapy is frequency dependent, HIF independent, and requires FOXO, PLCβ, Go/iα, IPMK, and IPPK.

---

Oxygen is a fundamental requirement for nearly all metazoans, however, environmental and physiological oxygen levels change and organisms must adapt rapidly in order to maintain cellular bioenergetics and sustain life processes. In terrestrial vertebrates, short term adaptation to limiting oxygen is accomplished by alterations in cardiopulminary rates, while long term adaptation is achieved by activation of the hypoxia controlled transcription factor HIF-1 (*1*). HIF-1 is highly conserved and plays similar roles in remodeling invertebrate physiology to meet the demands of hypoxic environments. In the nematode C. elegans, HIF-1 activation has been demonstrated to be required for embryonic survival in hypoxia, but is dispensable for suspended animation triggered by anoxia(*2*) Under normoxic conditions HIF-1 is produced constitutively and degraded by its negative regulators EGL-9 and VHL-1, but when oxygen concentrations decrease, HIF-1 is stabilized leading to activation of the genetic program to adapt to low oxygen(*2*). Several groups have shown that HIF-1 activation in normoxia, by inactivation of either EGL-9 or VHL-1, leads to an increase in stress resistance(*3*), reduced susceptibility to neurodegenerative disease, and increased lifespan(*4*). Interestingly, the increase in lifespan from genetic activation of HIF-1 is significantly larger than that observed when worms are cultured in hypoxic environments(*5, 6*). The reason for this difference is unclear, but may be due in part to the persistent physiological discordance between the abundance of atmospheric oxygen and the adaptive changes of the hypoxic genetic program, which leads to a large decrease in reproductive fitness but an increase in somatic stress resistance (*4*). This would explain why environmental hypoxia does not afford a similar increase in longevity as worms adapt to the environmental condition and maintain their reproductive output(*7*).

If adaptation to a fixed oxygen concentration underlies the relatively small effects of environmental hypoxia on *C. elegans* longevity, rapidly changing environmental oxygen would be predicted to decrease the ability of the worms to adapt and potentially increase lifespan. Such intermittent hypoxia regimes have previously been employed in athletic conditioning as well as preconditioning protocols frequently used prior to vascular surgery(*8, 9*). Indeed, previous work has shown that a brief hypoxic exposure during mid-life was sufficient to extend worm lifespan(*10*). We report here that lifelong intermittent hypoxia treatments (IHT) dramatically extend lifespan in *C. elegans* and that this effect is frequency and oxygen concentration dependent. We also show that mutations in the nematode homolog of the human inositol polyphosphate multikinase (IPMK-1) as well as the worm homologs of phospholipase C-beta4 (EGL-8), the trimeric G protein Go-**α** (GOA-1) and inositol pentakis phosphate kinase (IPPK-1) also suppress this lifespan extension.

## Results

### Whole life IHT doubles lifespan in wildtype worms

To test the lifelong effects of intermittent hypoxia exposure we constructed a computer-controlled oxygen delivery platform to modulate environmental oxygen over the course of a months long lifespan assay (Fig S1). We screened the effect of different oxygen concentrations on *C. elegans* lifespan and development. In general, all regimes with brief periods of room air led to large increases in lifespan (Fig 1a) coupled with developmental delays that were correlated with the duration of the hypoxic phase of the regime (Table S1). Interestingly, when the frequency of the intermittent hypoxia regimes were altered while maintaining the total number of moles of oxygen the animals were exposed to over their lifetime, we observed dramatically different results (Fig 1b). These data suggest that the acute response to oxygen deprivation plays a role in lifespan extension, rather than simply lowering the total number of moles of oxygen available for metabolism or oxidative damage over the course of the animal’s life. An equal 2hr IHT regime yielded the longest mean lifespan extension with the shortest developmental delay and was selected for further study. Animals reared in IHT are smaller (83% the size of their normoxic broodmates) and lay fewer eggs (63%) over an extended reproductive period compared to normoxic controls (Fig S2).

**Fig. 1.**
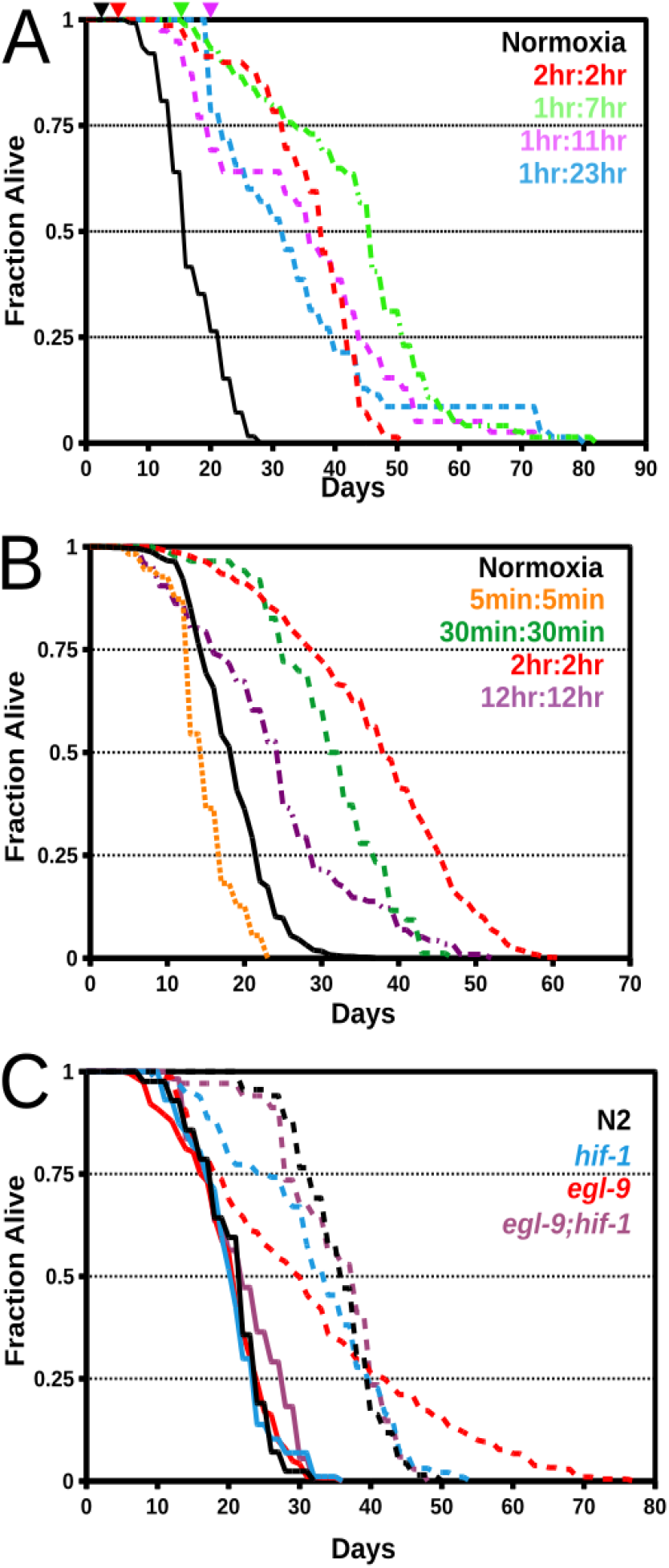
IHT doubles wildtype lifespan and does not require HIF-1 activity. A. Extended hypoxic phases increase lifespan but delay development, colored arrowheads indicate time of first egg lay. Normoxia treated survival in solid lines, IHT in dashed lines. Ratios normoxia:1000ppm oxygen. B. Hypoxia exposure frequency, not total lifetime oxygen exposure, determines longevity. C. HIF-1 is not required for IHT longevity. Statistics see Table S2.

### HIF-1 is not required for IHT induced longevity

We have shown previously that HIF-1 is required for the lifespan extension when grown in 5000ppm oxygen(*6*). Surprisingly, we find that HIF-1 is not required for the longevity of animals treated with IHT (Fig 1C). Loss of HIF-1 results in a subset of animals dying early in IHT, but the mean lifespan of the IHT population is still almost twice that of normoxic controls. Furthermore, constitutive stabilization of HIF-1 by mutation of EGL-9 has little effect on mean lifespan in IHT, although the overall shape of the survival curve and maximum lifespan is altered (Fig 1C). This effect is dependent on the activity of HIF-1, as an *egl-9;hif-1* double mutant does not show this phenotype (Fig 1C).

### IHT leads to genome-wide alterations of the transcriptome and activation of DAF-16 target genes

Because the canonical hypoxia response pathway is not required for the longevity of N2 animals in IHT, we performed RNAseq to determine the transcriptional changes during IHT. RNA was collected from wildtype worms raised in IHT on day 1 of adulthood; at both the end of the normoxic phase, and the end of the hypoxic phase. In addition to normoxic controls, libraries were also synthesized from *egl-9(jt307*) animals raised in IHT as well as N2 and *egl-9* animals grown in continuous 5000ppm oxygen. We observe that continuous hypoxia (5000ppm) activates many previously described HIF-1 target genes including *fmo-2* (*11*)(Fig S3a). Intriguingly, the transcriptional profile of IHT raised animals is distinct from both their normoxic broodmates and animals raised in continuous hypoxia, with only 39% and 36% of transcripts being increased or decreased respectively (Figs 2a,S4). Pathway enrichment analysis revealed an overrepresentation of several gene families including Beta fatty acid oxidation, sulfur amino acid metabolism and protein folding (Fig 2a, Table S3). RNAseq of *egl-9* animals raised in IHT reveals large changes in two components of the electron transport chain encoded by the mitochondrial genome (Fig S3b). CTC-1 the mitochondrially encoded subunit of cytochrome C oxidase is increased 10,000 fold compared to N2 animals in IHT and ATP-6, an integral membrane component of ATP synthase, is down regulated 3,000 fold. These enormous changes likely reflect an adaptive response to scavenge the decreased molecular oxygen for complex IV and potentially decrease ATP expended through reverse activity of ATP synthase, which can hydrolyze ATP and pump protons to maintain mitochondrial membrane potential during hypoxia(*12*).

**Fig. 2.**
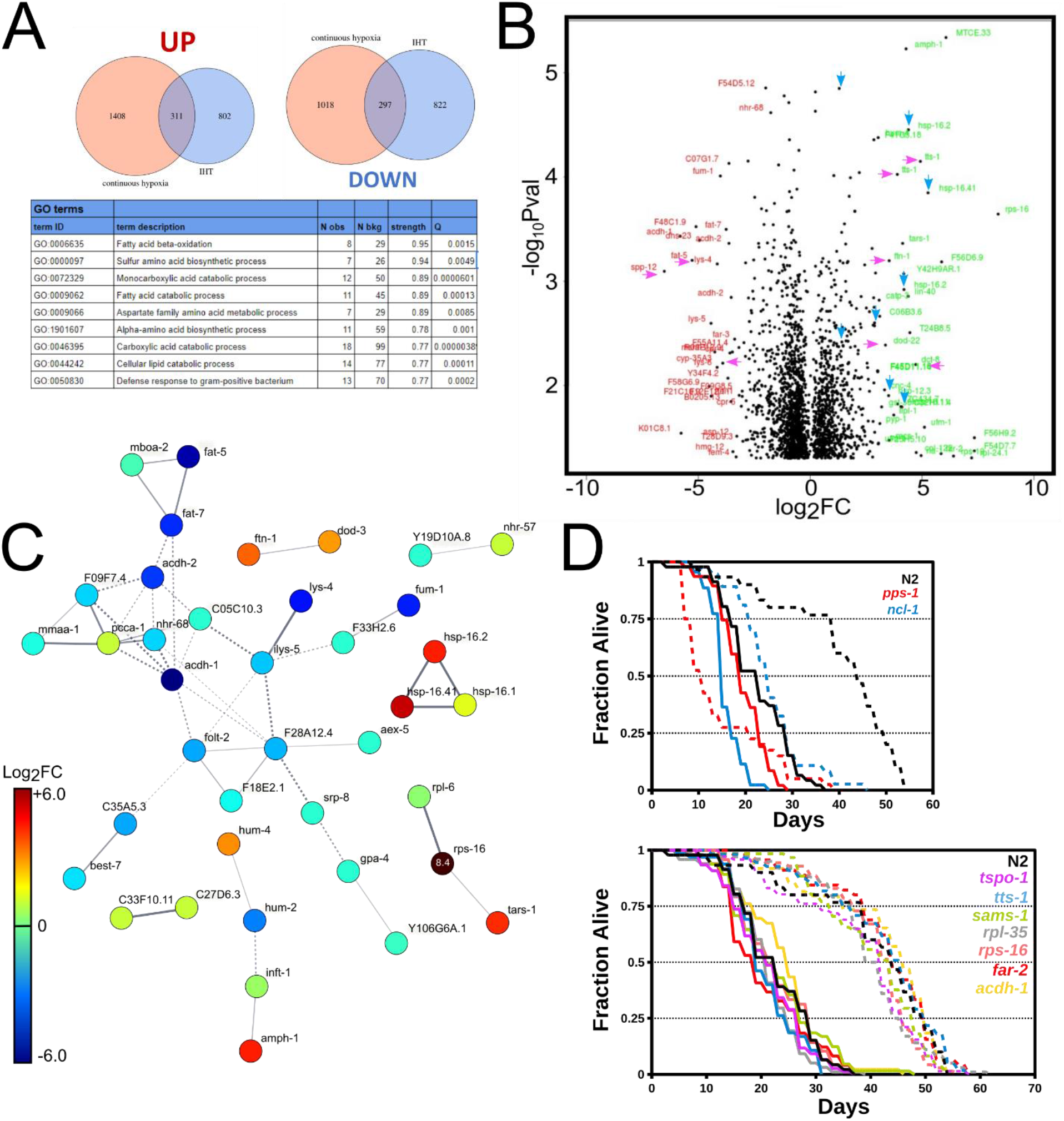
RNAseq analysis of animals in IHT. A. IHT is a distinct transcriptional state from continuous hypoxia (5000ppm oxygen). GO-term enrichment analysis of genes differentially regulated in IHT (Full list see Table S3). B. Volcano plot of transcript changes during the hypoxic phase of IHT animals compared to normoxic controls. Blue arrows indicate upregulated heat shock proteins, magenta arrows, a subset of genes previously shown to be regulated by DAF-16. C. Gene interaction network during hypoxic phase of IHT. Color indicates log2 of the fold change vs normoxic controls. D. Survival curves for genes involved in pathways identified via RNAseq. Normoxia treated survival in solid lines, IHT in dashed lines. Also see (Fig S6, Table S2).

Perhaps the best characterized longevity intervention in the worm is activation of the FOXO transcription factor DAF-16 via inactivating mutations in the IGF receptor DAF-2, which can double the lifespan of the worm(*13*). DAF-16 activity results in the transcription of numerous stress response genes and transcriptional profiling of *daf-2* animals has identified many genes required for their longevity(*14*). A subset (100) of the most highly differentially regulated genes by DAF-16 was disproportionately represented in our combined dataset of IHT regulated genes (26%, representation factor 6.8, P= 3.9×10^-15^). Heat shock proteins (HSPs), under control of the HSF-1 heat shock transcription factor, function along with DAF-16 to promote longevity in daf-2 animals(*15*), and expression of an HSP transgene is predictive of longevity(*15–17*). In our IHT samples we observe 7 different HSPs upregulated more than 1.5 fold over normoxic controls(Fig 2B).

In addition to the large number of transcriptional changes observed between normoxia and IHT, there were multiple genes that showed dynamic changes during the phases of IHT exposure (Fig S5). R102.4, which encodes the worm L-allothreonine aldolase, was the most significantly upregulated gene between the IHT phases. Homologs of this enzyme convert L-allothreonine to glycine and acetaldehyde, which has been shown to protect anoxic cells from reductive stress(*18*). Intriguingly, we also observe that TARS-1, the threonyl amino-acyl tRNA synthetase, was also highly upregulated in IHT (Fig 2c). Forward genetic screens identified *tars-1* loss of function mutants as being protected from lethal hypoxia, through a mechanism thought to involve global repression of translation(*19*). We also observe large changes in several other genes also involved in the translation machinery, including the ribosomal protein gene *rpl-35*, misregulation of which has been implicated in human cancer(*20*). RPL-35 and RPS-16 RNAi animals showed a profound developmental delay in both normoxia and IHT, however longevity in IHT was unchanged from wildtype (Fig 2d), suggesting that the observed downregulation of these ribosomal proteins may underlie the developmental delay observed in IHT but not its effect on longevity. Control of ribosome biogenesis and protein synthesis has previously been shown to influence both longevity and survival during hypoxia(*21–23*). Loss of function mutations in the worm homolog of the tumor suppressor BRAT/TRIM3, *ncl-1*, which has been shown to suppress hypoxia survival and several longevity interventions(*19, 23*), resulted in short lived animals that showed a partial suppression of IHT longevity (Fig 2d). Also differentially regulated between the two phases was the worm homolog of the yeast mitochondrial adenosine 5’-phosphosulfate (APS) and 3’-phospho-adenosine-5’-phosphosulfate PAPS transporter MRX21(F55A11.4). While RNAi of F55A11.4 had no effect on IHT survival (Fig S6), RNAi of the worm PAPS synthase, PPS-1, completely suppressed IHT longevity by rendering the worms extremely sensitive to hypoxia but only reducing normoxic lifespan slightly (Fig 2D). While we do not yet know the mechanism by which loss of PPS-1 suppresses IHT longevity, it is notable that in dissimilatory sulfate reducing (DSR) bacteria cellular energy is produced via electron-transport based phosphorylation in the absence of molecular oxygen using PAPS as an electron acceptor. While DSR has not been described in animals, assimilatory sulfate reduction is critical to sulfur amino acid metabolism and worms excrete hydrogen sulfide during nutritive stress(*24, 25*). Taken together, while some single gene knockouts produced significant changes to normoxic lifespan or the IHT response(Fig 2D,S6), none of the single gene knockouts tested restored normoxic lifespan to IHT treated animals or resulted in an equivalent lifespan extension, suggesting that multiple pathways and transcriptional changes underlay the response to IHT.

### The FOXO transcription factor DAF-16 is required for IHT longevity

Because of the large number of DAF-16 target genes upregulated in IHT, we tested whether mutations in *daf-2* and *daf-16* affected the IHT phenotype. Although IHT was able to extend the lifespan of *daf-16*(*mu86*) animals, the magnitude of lifespan extension was significantly reduced relative to the effect of IHT in wild type. Interestingly, IHT failed to significantly extend the median lifespan of *daf-2*(*e1370*) animals but resulted in a large extension of maximum lifespan (Fig 3a). We previously observed that exposure to hypoxia caused a DAF-16::GFP fusion protein to rapidly accumulate in the nucleus(*6*). Consistent with this, we observe that when placed into 1000ppm oxygen, 100% of TJ356 animals have nuclear accumulation of DAF-16 within 50 minutes(Fig 3b). This time delay may explain why IHT regimes with shorter hypoxic phases were less effective (Fig 1a). In order to determine what factors triggered DAF-16 nuclear localization in IHT, we performed a temperature sensitive forward genetic screen in the TJ356 background to identify animals with altered kinetics of DAF-16::GFP localization in hypoxia. Of the hundreds of F2 lines we isolated, 16 showed stable alteration of DAF-16 nuclear localization in hypoxia and we named these strains *hyds*, for hypoxia daf-16 suppressors. One of the most penetrant of these strains *hyds-2*, was backcrossed to TJ356 and then outcrossed to N2. In addition to slow DAF-16::GFP nuclear localization in hypoxia, *hyds-2* worms show a temperature sensitive growth arrest phenotype at 26.5C, everted vulva, and germline defects (Fig S7). Like *daf-16* animals, *hyds-2* animals showed a partial suppression of the IHT longevity phenotype (Fig 3c).

**Figure 3.**
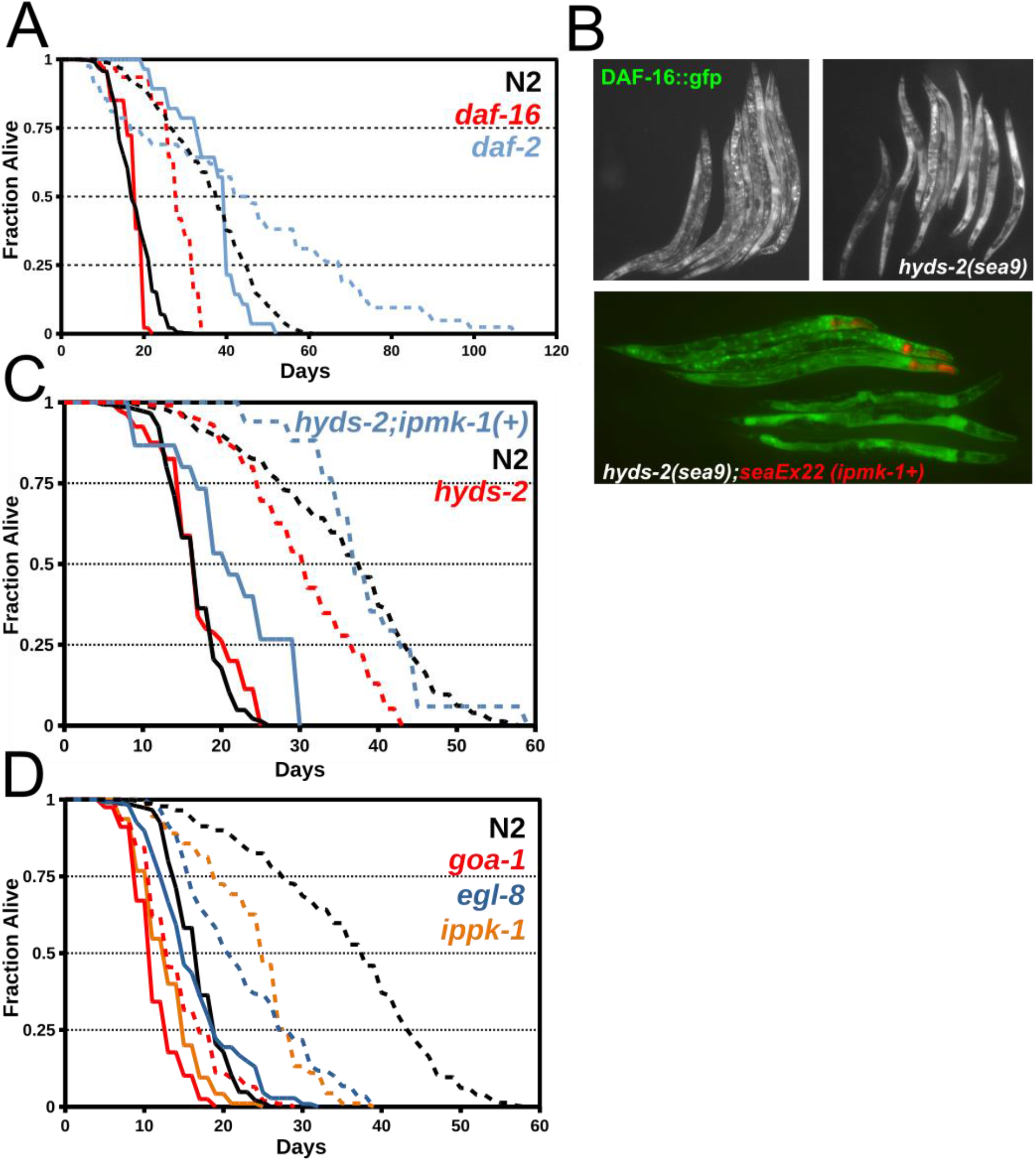
Insulin IGF signaling is required for IHT and regulators of inositol signaling pathways suppress IHT longevity. Normoxia treated survival in solid lines, IHT in dashed lines. A. Survival curves for *daf-2(e1370*) and *daf-16(mu86*) animals in IHT. B. DAF-16::GFP localization in day 1 adults following a 24hr exposure to 25C and 50 min in 1000ppm O2. Clockwise TJ356 strain, TJ356 strain carrying the *hyds-2(sea9*) allele, *hyds-2(sea9*) animals rescued with wildtype copy of *ipmk-1 (seaEx22*) marked with myo-2::mCherry(top), co-treated broodmates lacking transgene (bottom). C. Survival curves of *hyds-2* and *hyds-2;ipmk-1+* rescue animals. D. Survival curves of other previously described modifiers of inositol phosphate signaling pathways also required for full IHT longevity(Table S2).

To identify the mutated gene(s), outcrossed *hyds-2* animals were crossed to the Hawaiian strain and F2 hybrids were pooled, sequenced, and analyzed with the CloudMap software package(*26*). Strong disequilibrium of Hawaiian SNPs was observed on linkage group IV and several severe loss of function mutations were observed in the linked region, including a splice site mutation leading to a premature stop in the worm homolog of the human enzyme inositol polyphosphate multikinase (Fig S8)(ZK795.1). Injection of a 9kb fragment of the wildtype ZK795.1 gene rescued the *hyds-2(sea9*) phenotypes (Fig 3b) and we renamed the strain *ipmk-1(sea9*). Rescue of the IPMK-1 mutation in *hyds-2(sea9*) with a wildtype copy of *ipmk-1* also restored the IHT lifespan effects to wildtype levels (Fig 3c). We created an *ipmk-1* reporter transgene with a 3kb promoter region of the *ipmk-1* gene cloned upstream of mCherry and injected it along with a histone GFP (PCFJ420) into wildtype animals. IPMK-1 was first expressed during embryogenesis in the E lineage (Fig S9), with expression spreading to many other tissues in larvae with the highest levels of expression seen in a subset of sensory neurons and the spermathecal precursor SS cell(Fig S9). IPMK-1 and its inositol polyphosphate products have been shown to interact with several pathways known to influence the rate of aging including mTOR, AMPK, and insulin IGF(*27*). We hypothesized that other components or regulators of inositol signaling pathways may also regulate IHT longevity. Loss of function mutations in the worm inositol pentakisphosphate 2-kinase IPPK-1(*28*), phospholipase C beta 4 homolog EGL-8(*29*), and the Gi/o-protein GOA-1(*30*) all blocked the longevity of IHT to an even greater extent than loss of IPMK-1 (Figure 3d). In particular, loss of GOA-1 resulted in a nearly complete abrogation of IHT lifespan extension. GOA-1 is expressed throughout the nervous system and several other tissues including the spermatheca, where we observe the highest levels of IPMK-1 expression.

The protective effects ischemic preconditioning (IPC) in mammalian systems requires the activity of G-protein coupled receptors, and inhibition of Gi/o with pertussis toxin can abrogate this protection(*31*). While IPC has been studied for decades, and the use of various IHT regimes to improve athletic performance equally well established(*32*), the clinical use of hypoxia in western medicine has been relatively limited(*33*). However, recent work shows that exposure to continuous hypoxia can rescue mitochondrial dysfunction in a mouse model of the mitochondrial disease Leigh syndrome, while an IHT regime failed to show a similar results(*34, 35*). Mitochondria are the primary consumers of molecular oxygen in the cell and mitochondrial dysfunction is one of the key hallmarks of aging(*36*). The role of oxygen in the aging process, primarily as a putative source of damaging ROS has been hypothesized for decades, however low oxygen therapies have shown limited results in increasing longevity(*37*). Our results suggest that organismal responses to low oxygen are complex, and that sensory or reproductive signals triggered by acute hypoxia exposure, rather than decreased total lifetime oxygen levels, can trigger endogenous protective pathways to dramatically increase lifespan in wildtype animals. Further studies are warranted to determine whether the effects of IHT on longevity are conserved in mammals with the potential to promote healthspan and lifespan in humans.

## Acknowledgments

Some strains were provided by the CGC, which is funded by NIH Office of Research Infrastructure Programs (P40 OD010440). Stan Fields, Josh Cuperus, Scott R. Kennedy, and Mary-Claire King for use of equipment for RNAseq experiments. James Priess, Alexander Mendenhall, Karre Fisher, Brandon Berry, Shane Rae, Alan Herr, and Michael Ailion for discussions, worm strains, and plasmids.

## Author contributions

Conceptualization: JNP, MBR, MK

Methodology: JNP, MBR, MK

Investigation: JNP, EC, KMP, MC, CT, JT, KH, AV, MLA, NA, SCL

Visualization: JNP

Funding acquisition: JNP, MBR, MK

Project administration: JNP,MBR,MK

Supervision: JNP,MBR,MK

Writing – original draft: JNP

Writing – review & editing: JNP,MK

## Data and materials availability

All data, code, strains and materials used in the analysis are available at github.com/jasonnpitt/medea, the CGC, NCBI SRA, or by request from the corresponding authors.

## Materials and Methods

### Nematode Culture and lifespan analyses

Worms stocks were grown on standard NGM media with an E. Coli OP50 food source. For manual lifespan experiments NGM plates were supplemented with antibiotics and IPTG and fed bacteria expressing dsRNA for the maternal effect lethal *pos-1* gene to prevent offspring hatching as has previously been described, rather than using FUDR(*38*). To control oxygen levels, NGM plates were placed in sealed chambers (Mitsubishi rectangular jars, Mitsubishi Gas Chemical, Tokyo, Japan) fitted with 1/16 inch barbed luer gas ports (MasterFlex, Gelsenkirchen, Germany) and supplied with humidified hypoxic gas mixtures controlled by the system described in the supplemental materials (Supp Fig1). Specified oxygen concentrations were generated using mass flow controllers (Sierra Instruments, Monterey, CA; Aalborg Instruments, Orangeburg, New York) to mix nitrogen and room air supplied by an air compressor (California air tools, Torrance, California) and the oxygen concentrations verified with a paramagnetic oxygen analyzer (Sybron|Taylor, ServoMex, Milwaukee, Wisconsin) spanned using supplied hypoxic gas standards (0ppm, 1000ppm, 5000ppm, 50,000ppm O2, Praxair, Seattle, WA). All experiments were performed in an iso-therm incubator (Torey Pines Scientific, Carlsbad, California) at 20C unless otherwise noted. All lifespans were started from synchronized worm populations generated by hypochlorite treatment (*39*) from a population of gravid adult worms and allowing their eggs to hatch overnight in 6mL of M9 at 20C in an empty 60mm petri dish, before being pipetted onto *pos-1* RNAi plates and being placed into the various atmospheres. Worms were removed from the chambers and checked every other day manually during the normoxic phase of the experiment for regimes that included hypoxia. Worms that appeared unmoving were touched with a platinum wire and worms that did not move within 10 seconds of stimulus were removed from the plate and recorded as dead. Worms that were ruptured were not censored but marked as dead when their evoked movements ceased. WormBot RNAi experiments were performed as previously described (*40*), animals were treated as above but grown from hatching to L4 in the presence of the bacteria expressing the dsRNA of interested on 60mm NGM plates supplemented with antibiotics, IPTG, and nystatin. When animals reached L4 stage they were transferred to sealed 12-well plates modified with 1/16 inch barbed luer gas ports containing RNAi media + FUDR to prevent hatching. Lifespan curves and statistics were calculated using OASIS2 (*41*). Strain genotypes listed in Table S2.

### DAF-16::GFP localization in hypoxia

Nuclear translocation of the DAF-16::GFP transgene (*42*) in hypoxia treated animals was observed using a Zeiss Lumar.v12 stereo fluorescence microscope (Carl Zeiss AG, Oberkochen, Germany). Animals were placed on standard seeded NGM plates in 400mL Mitsubishi chambers (see above) or 15cm petri dish chambers fitted with lids modified with gas ports and sealed with petroleum jelly. 1000ppm oxygen balanced nitrogen flowed into a bubble flask containing water for humidification and then into the chamber at 1 chamber volume per minute controlled via a rotometer. Animals that had any visible nuclear DAF-16::GFP enrichment in the intestinal nuclei were scored as positive. For micrographs, plates were removed from the atmosphere and immediately inverted over a 500mL jar of iodine crystals for 20 seconds to fix the worms in iodine vapor. Animals were then immediately grouped into position with a worm pick and photographed using a Zeiss Axiocam MRm camera and Axiovision software.

### RNAseq library prep, sequencing and analysis

For RNA seq experiments ~600 animals were grown on 10cm plates kept in their respective atmospheres at 20C until the worms reached day 1 of adulthood. In order to isolate the animals for RNA purification within 90 seconds of removal from the atmosphere we developed a protocol for rapidly purifying the worms from the bacterial food source using a 40um FlowMi pipette tip filter (Bel-Art, Wayne, NJ). Briefly, the NGM plate was removed from the atmosphere and immediately flooded with 1700uL M9 with 0.01% Tween20. The plate was tilted so that the worms collected in the bottom corner of the plate and were aspirated into a 1000uL pipette tip. With the worms still in the pipette tip, a 40um FlowMi filter was added to the end of the pipette tip. The M9 containing bacterial contaminants was expelled through the filter retaining the worms in the pipette tip. The outside of the filter was washed in a stream of deionized water and 1000uL of fresh M9 /w tween drawn into pipette tip. This wash cycle was repeated rapidly 4 times. Following the final wash ~200uL of M9 w/ tween followed by ~200 uL air was drawn into the pipette tip bringing the worms out of the filter. The FlowMi tip was removed and the washed worms pipetted into 500uL of Trizol in an eppendorf tube, vortexed briefly and then immediately flash frozen on liquid nitrogen. Worms were then stored at −80C until all samples had been collected. All samples were prepared in biological triplicate. Samples thawed and then disrupted with an eppendorph tube pestle and total RNA purified using a Direct-zol plus RNA miniprep kit (Zymo Research, Irvine, CA) following the manufacturer’s instructions. Total RNA was quantified by UV spectrophotometry (Nanodrop, Thermo Scientific, Waltham, MA) and RNAseq libraries constructed using an NEBnext Ultra RNA library prep kit with a poly A purification module (New England Biolabs, Ipswich, MA). Libraries were indexed using multiplex adapters made by IDT (Coralville, IA) and quantified using a Tapestation 4200 (Agilent Technologies, Santa Clara, CA) with D1000 tapes and a Kappa Illumina Library Quantifcation kit (Kapa Biosystems, Wilmington, MA) using an AriaMX realtime PCR machine (Agilent Technologies, Santa Clara, CA). Quantified libraries were pooled and sequenced on an Illumina NextSeq using a high capacity 75 read flow cell (Illumina, San Diego, CA) following the manufacturer’s instructions. Data was processed using the HISAT2/StringTie/Ballgown pipeline(*43*) and figures generated in R.

### Temperature Sensitive Forward Genetic Screening and Genetic Mapping

The TJ356 strain was synchronized by bleaching and 5000 L4 animals placed into 4 mL M9 with 20uL EMS (Sigma-Aldrich, St. Louis, MO) for 4 hours at room temperature. P0 animals were transferred to 15C and grown for 48 hours and then bleached to obtain a synchronized population of F1 animals, which was split into 4 groups of 10,000 animals and grown until gravid at 15C. F1 animals were then bleached to obtain synchronized populations of F2 animals which were grown at 15C until the L4 larval stage when they were upshifted to 25C for 24 hours. Animals were plated at a density of 5000 animals per 10cm NGM plate and then placed into 1000ppm oxygen balanced nitrogen for 1 hour and then screened for DAF-16 activation as described above. Mutants were cloned to individual 35mm plates and returned to normoxia at 15C. Strains were passaged and rescreened for several generations to ensure homozygosity and given the designation *hyds* (***hy***poxia ***d***af-16 **s**uppressors). *hyds-2* mutants were backcrossed 4 times to the original TJ356 strain and then crossed to the Hawaiian isolate CB4856 and mapped using RFLP and WGS of recombinant F2 lines as described previously(*26*). Generation of fertile males of the TJ356 roller strain was accomplished by feeding roller worms RNAi for the *rol-6* gene (J.R. Priess, personal communication).

### IPMK-1 Transgene Construction and Rescue

To identify the molecular lesion in the *hyds-2* strain a 9KB PCR fragment of the ZK795.1 gene was cloned into the PUC19 plasmid by Gibson assembly (New England Biolabs, Ipswich, MA). The *seaEx22* plasmid along with a myo-2:mCherry (PCFJ90) co-injection marker was injected into a *hyds-2* strain that had been backcrossed 4 times to TJ356 and stable transgenic lines selected. A fluorescent transcriptional reporter for IPMK-1 was produced by cloning a 3kb fragment taken from the upstream region of the ZK795.1 using a PCFJ90 backbone using Gibson assembly.

