## Supporting Online Information for "Intermittent hypoxia therapy engages multiple longevity pathways to double lifespan in *C.elegans*"

**Supplementary Information**

A.

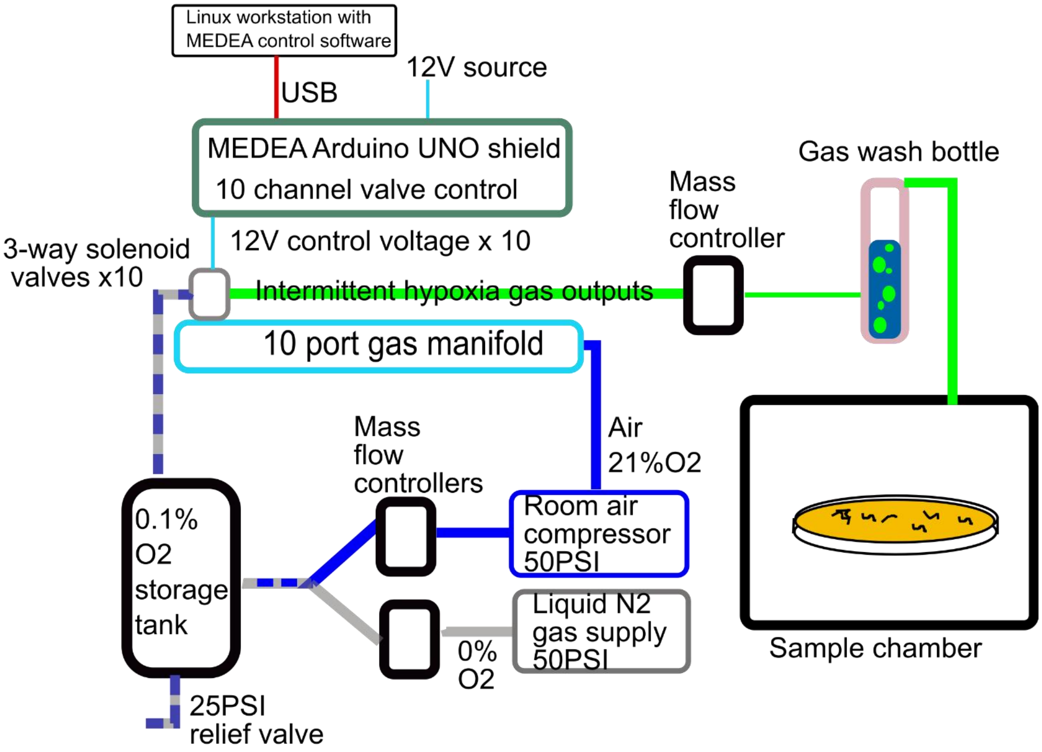

B.

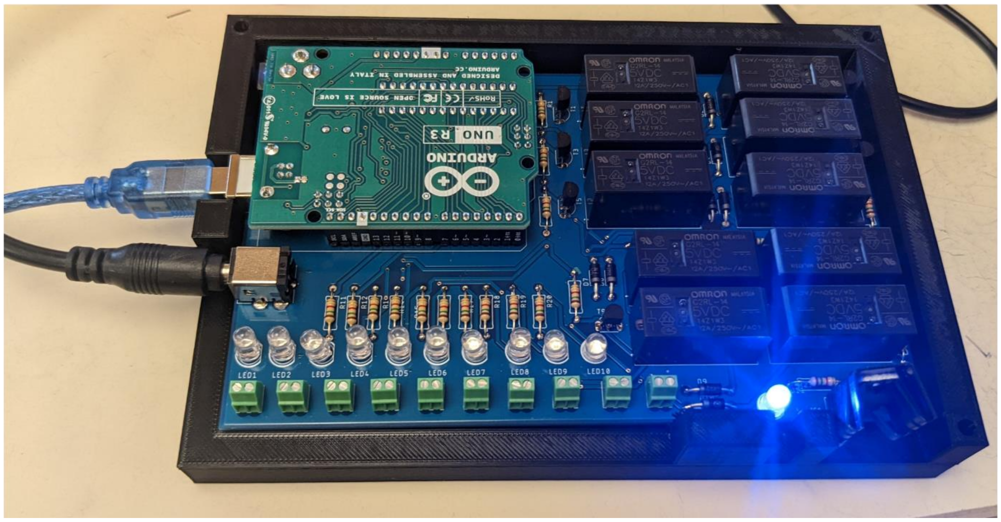

**Figure S1. Gas control system for IHT experiments.** A. Schematic illustration of the gas sources and flow control. For simplicity only one output channel of the 10 channel system is shown. B. Custom Arduino shield for 10 channel computer gas control. Control software, firmware, BOM, and PCB files available at http://github.com/jasonnpitt/medea

| Normoxic phase Duration | 1000ppm $O_2$ Phase Duration | Days to adulthood (days) | Developmental delay (days) |
|---|---|---|---|
| 24hr | 0 | 2 | 0 |
| 2hr | 2hr | 5 | 3 |
| 1hr | 7hr | 15 | 13 |
| 1hr | 11hr | 20 | 18 |
| 1hr | 23hr | NA | $\infty$ |

**Table S1. Developmental delays observed in IHT at 20C.** Longer hypoxic exposures resulted in longer lifespan extensions however a 2hr -2hr cycle resulted in doubling of normoxic lifespan with only a 3 day developmental delay.

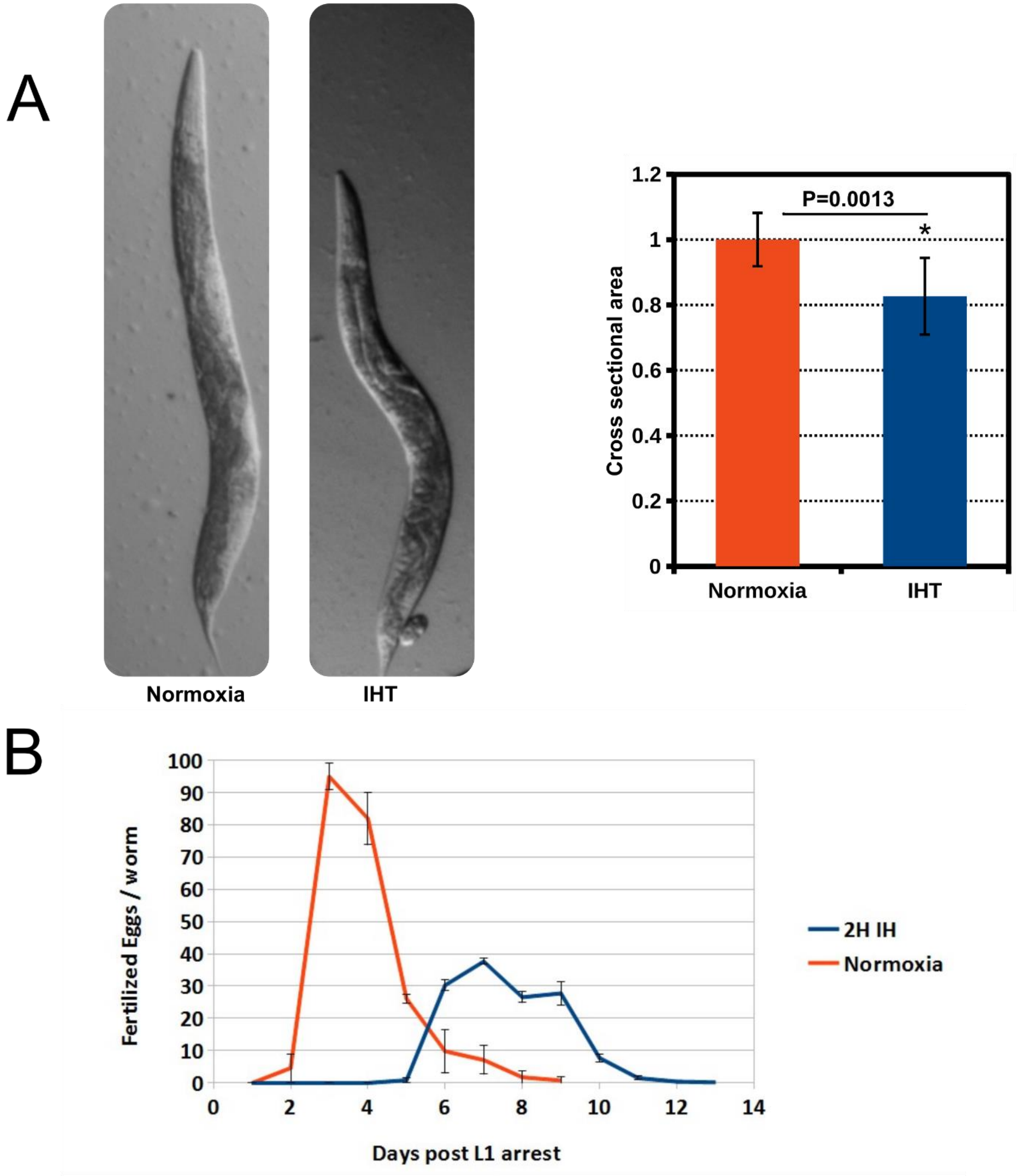

**Figure S2. IHT decreases adult body size and reduces fecundity**. A. IHT results in small adults (83% relative cross-sectional area). B. Brood size is decreased (63% of normoxic controls) and reproductive period extended in IHT animals.

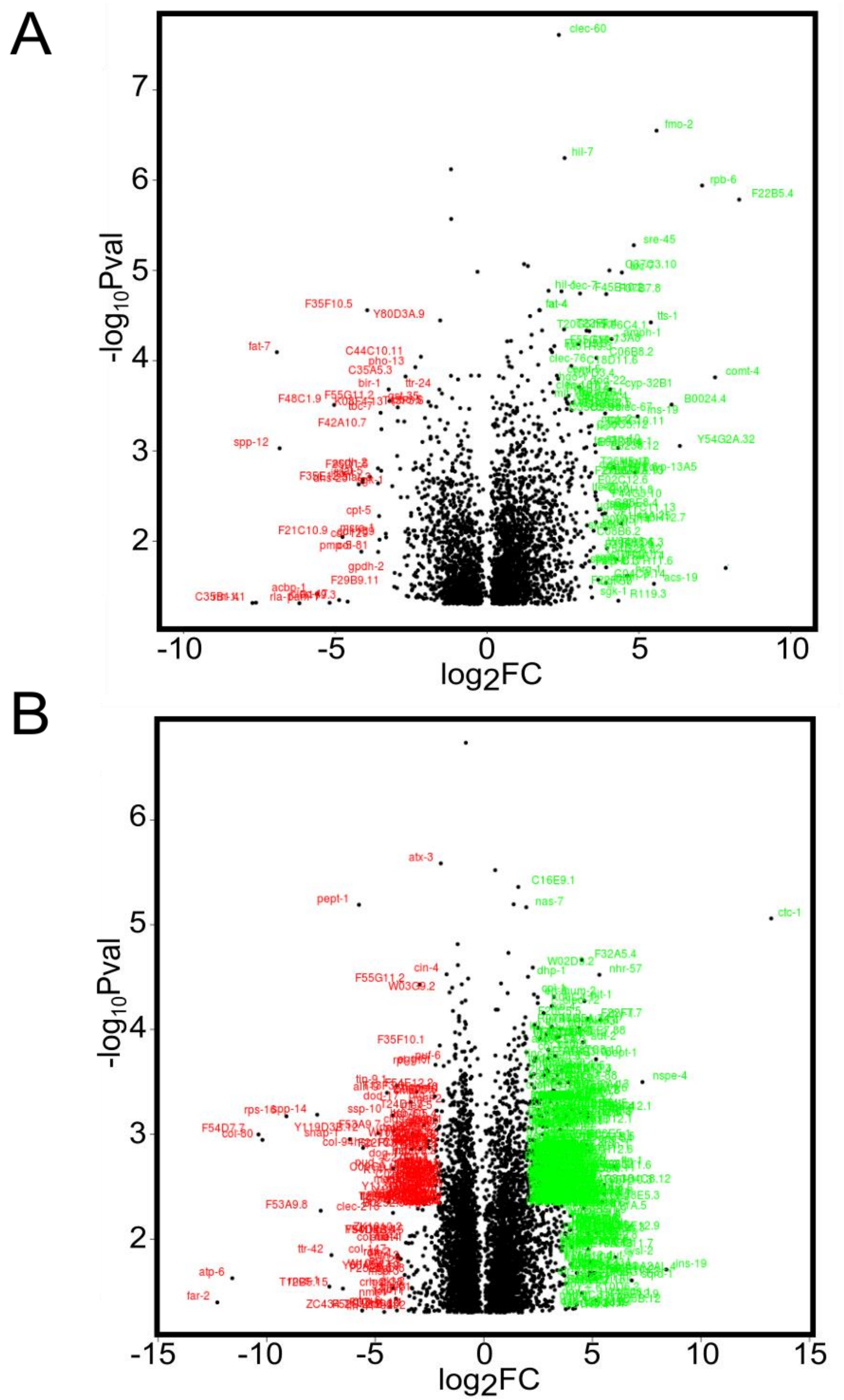

**Fig S3. Volcano plots of continuous hypoxia and *egl-9(jt307)* in IHT.**

A. N2 day 1 adults raised in continuous 5000ppm $O_2$ vs N2 normoxia. B. *egl-9(jt307)* IHT vs N2 IHT

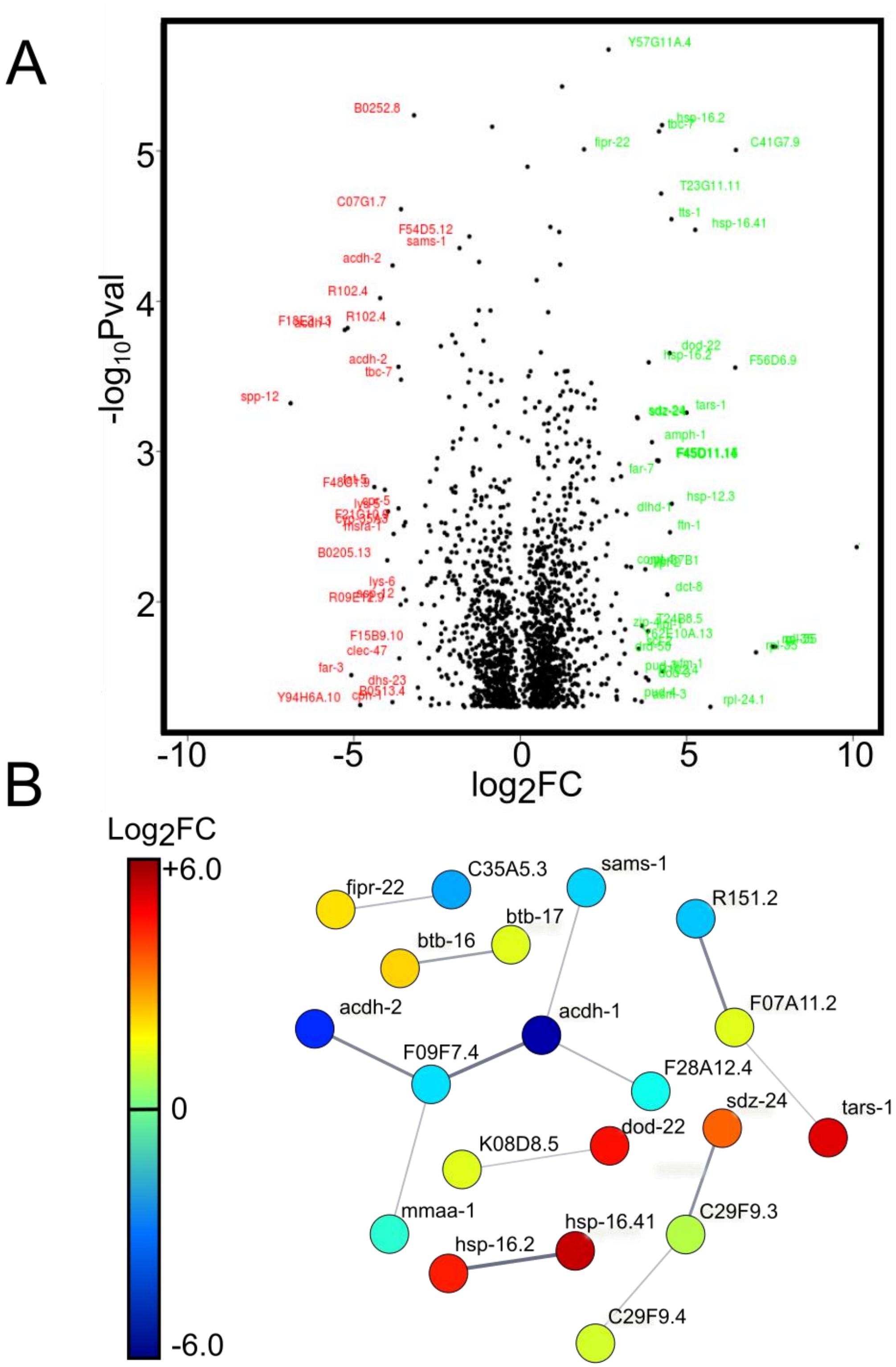

**Figure S4. Transcript changes observed in normoxic phase of IHT treated animals compared to normoxic controls.** A. Volcano plot of the most highly differentially regulated genes. B. Gene interaction network in normoxic phase generated by STRING(string-db.org)(also see Fig 2).

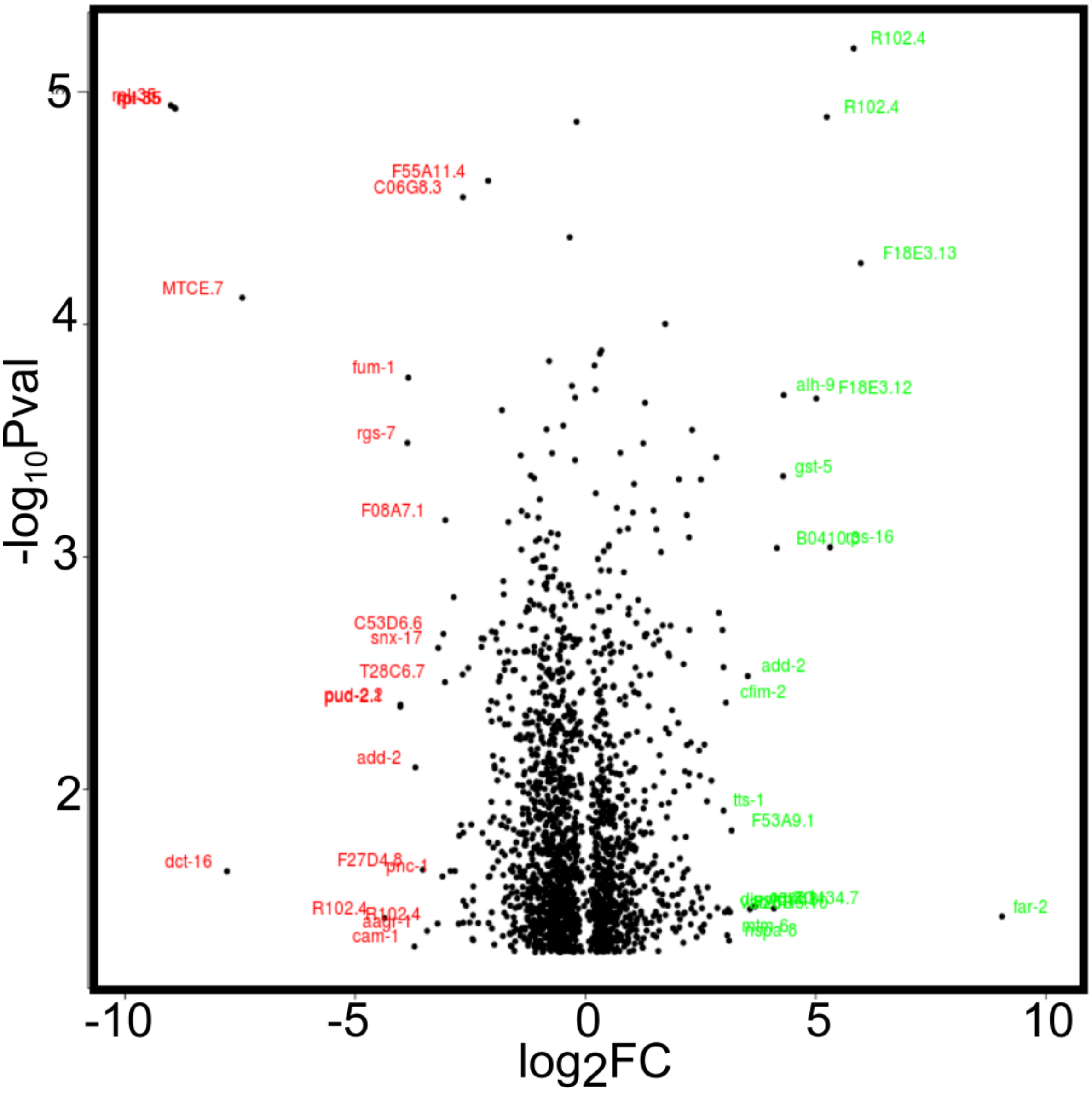

**Figure S5. A smaller subset of transcripts change expression during IHT cycling.** Volcano plot of transcript changes observed in hypoxic phase of IHT treated animals compared to normoxic phase of IHT treated animals.

A

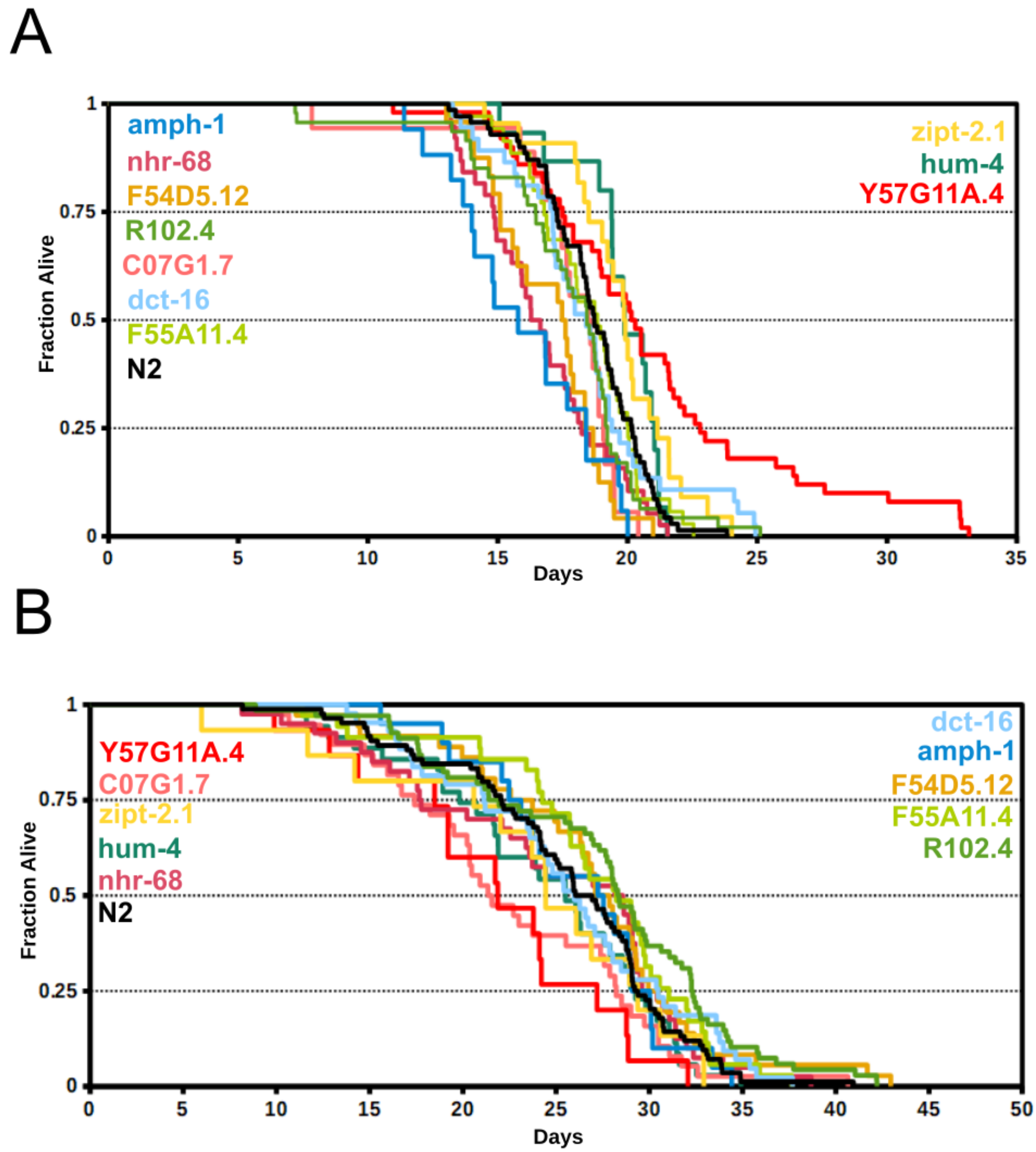

**Figure S6. WormBot generated survival curves of RNAi treated animals in IHT.** A. IHT raised animals transferred to normoxic WormBot plates as L4s B. Whole life IHT animals transferred to IHT WormBot plates as L4s. Gene names ranked in order of mean lifespan for each treatment. For statistics see table S2.

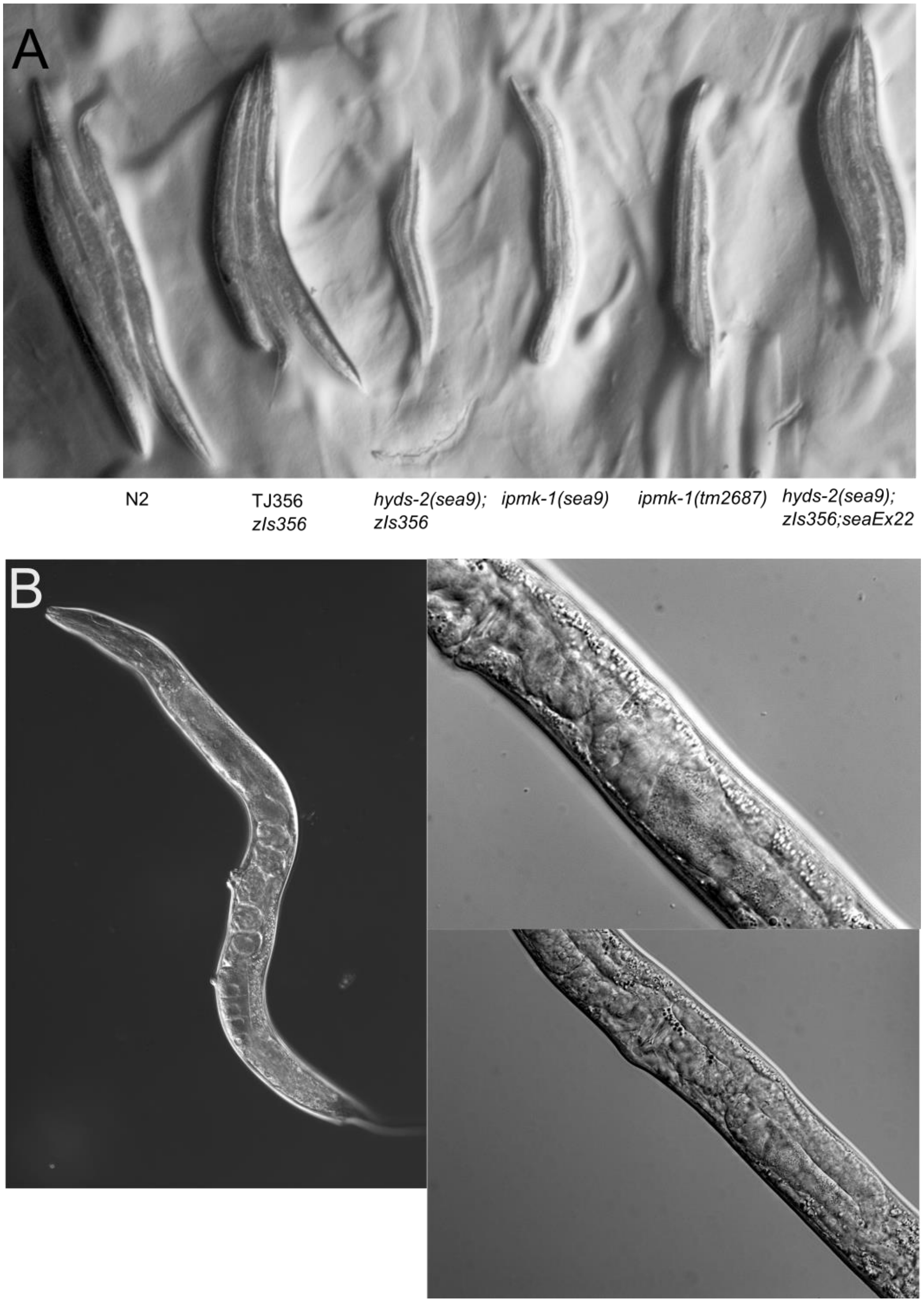

**Figure S7. *hyds-2(sea9);zIs356* phenotypes.** A. *hyds-2* and control animals grown from egg at 26.5C, *ipmk-1* loss of function leads to a variable lethal developmental arrest. This phenotype is rescued by wildtype IPMK-1 expression (*seaEx22*). B. DIC images of vulval and germline phenotypes. Left: pvl *hyds-2(sea9);zIs356* animal raised at permissive temperature (15C). Right: underdeveloped and disorganized germlines in *ipmk-1(sea9)* animals raised at 26.5C.

A

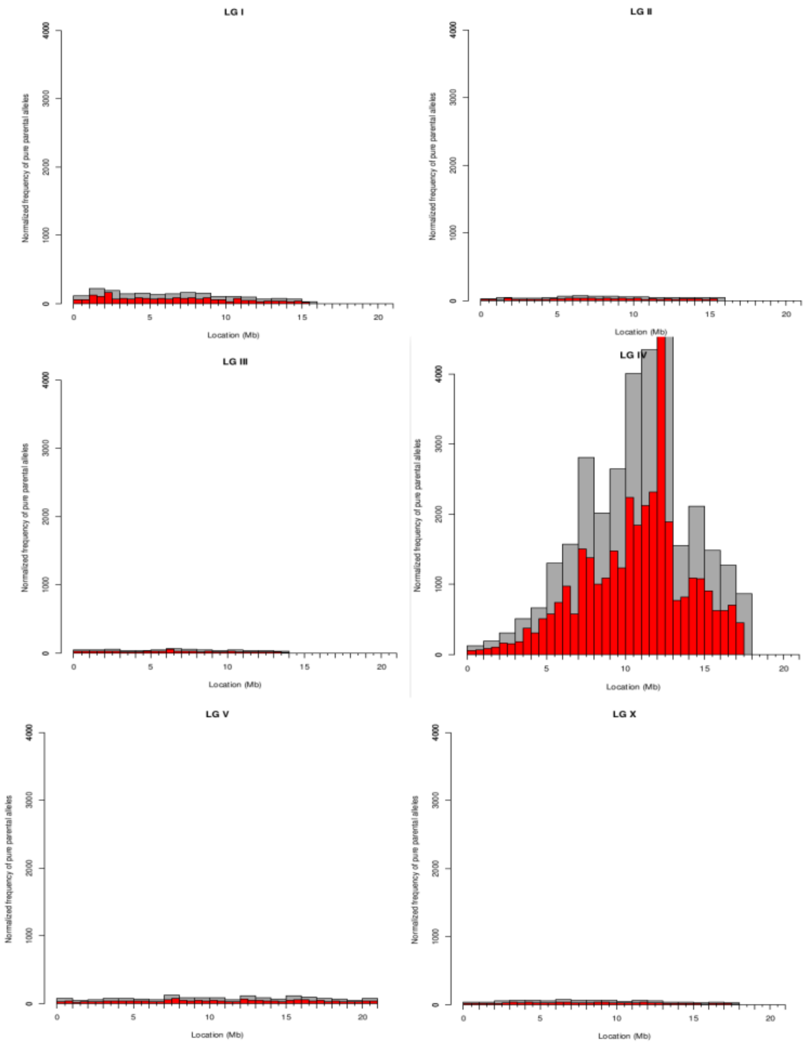

B

| | | | | | | | | | | | | | | | | | | |
|---|---|---|---|---|---|---|---|---|---|---|---|---|---|---|---|---|---|---|
| IV | 11226249 | A | C | SNP | Hom | 188.39 | 8 | | C29F4.1 | col-125 | protein_cod | C29F4.1 | exon_IV_112 | | 2 | NON_SYNO | Q/P | cAa/cC |
| IV | 11616803 | A | C | SNP | Hom | 50.96 | 6 | | Y24F12A.4 | Y24F12A.4 | pseudogene | Y24F12A.4 | | | | TRANSCRIPT: Y24F12A.4 | | |
| IV | 11654592 | C | T | SNP | Hom | 868.05 | 24 | | ZK809.4 | ent-1 | protein_cod | ZK809.4 | exon_IV_116 | | 4 | NON_SYNO | G/E | gGa/g |
| IV | 11725149 | C | T | SNP | Hom | 701.24 | 20 | | K01A6.2 | magi-1 | protein_cod | K01A6.2d | exon_IV_117 | | 4 | NON_SYNO | P/S | Cct/Tc |
| IV | 11725149 | C | T | SNP | Hom | 701.24 | 20 | | K01A6.2 | magi-1 | protein_cod | K01A6.2c | exon_IV_117 | | 10 | NON_SYNO | P/S | Cct/Tc |
| IV | 11725149 | C | T | SNP | Hom | 701.24 | 20 | | K01A6.2 | magi-1 | protein_cod | K01A6.2b | exon_IV_117 | | 5 | NON_SYNO | P/S | Cct/Tc |
| IV | 11725149 | C | T | SNP | Hom | 701.24 | 20 | | K01A6.2 | magi-1 | protein_cod | K01A6.2a | exon_IV_117 | | 10 | NON_SYNO | P/S | Cct/Tc |
| IV | 11725149 | C | T | SNP | Hom | 701.24 | 20 | | K01A6.2 | magi-1 | protein_cod | K01A6.2e | exon_IV_117 | | 3 | NON_SYNO | P/S | Cct/Tc |
| IV | 12147659 | C | T | SNP | Hom | 196.65 | 9 | | B0001.2 | B0001.2 | protein_cod | B0001.2 | exon_IV_121 | | 9 | NON_SYNO | E/K | Gaa/A |
| IV | 12439159 | T | C | SNP | Hom | 750.11 | 20 | | C25G4.1 | clec-185 | protein_cod | C25G4.1 | | | | SPLICE_SITE_DONOR | | |
| IV | 12514485 | T | G | SNP | Hom | 47.39 | 2 | | F53B2.7 | F53B2.7 | pseudogene | F53B2.7 | | | | TRANSCRIPT: F53B2.7 | | |
| IV | 12550646 | C | T | SNP | Hom | 471.72 | 13 | | ZK795.1 | ZK795.1 | protein_cod | ZK795.1 | exon_IV_125 | | 4 | STOP_GAIN | Q/* | Caa/Ta |
| IV | 12878236 | C | T | SNP | Hom | 124.77 | 4 | | ZK896.5 | ZK896.5 | protein_cod | ZK896.5.2 | exon_IV_128 | | 6 | NON_SYNO | G/S | Ggc/A |
| IV | 12878236 | C | T | SNP | Hom | 124.77 | 4 | | ZK896.5 | ZK896.5 | protein_cod | ZK896.5.1 | exon_IV_128 | | 6 | NON_SYNO | G/S | Ggc/A |

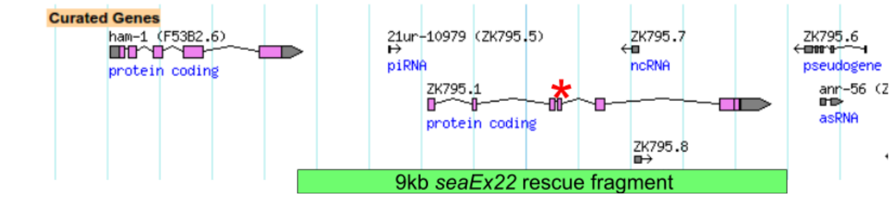

**Figure S8. Genetic mapping of *hyds-2(sea9)* mutation.** A. Linkage disequilibrium across *C. elegans* genome in recombinant lines. B. Genetic lesions observed in the linked area and genomic diagram of position of *ipmk-1(sea9)* allele location (Q108*) and *ipmk-1* rescue construct *seaEx22*.

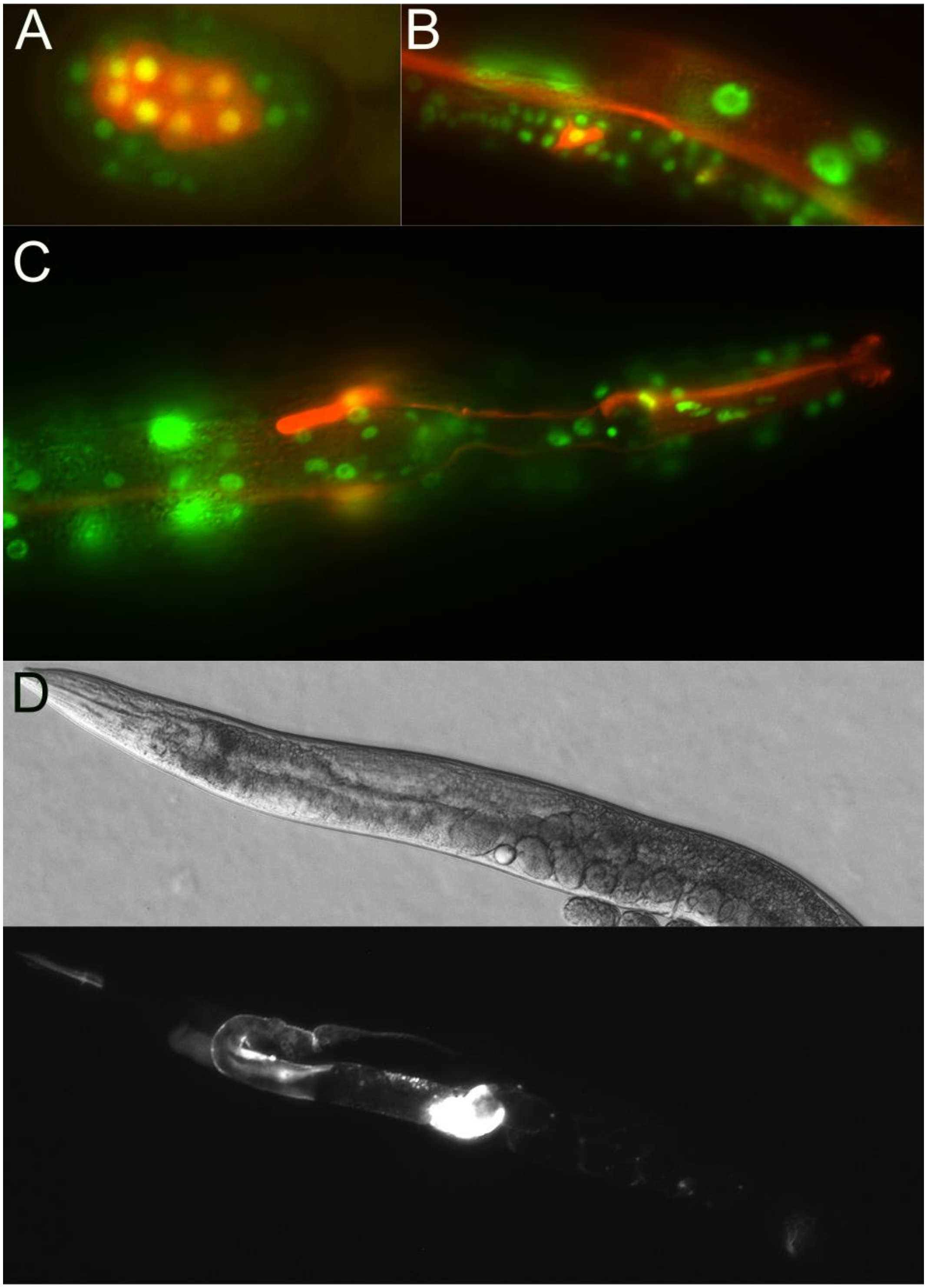

**Figure S9. Expression of IPMK-1 transcriptional reporter.** A. IPMK-1 expression is first detected in the embryonic intestinal precursors at the E8 cell stage. B. Low levels of expression are detected in many larval tissues but highest levels seen in the spermathecal precursor cells at the early L3 stage. C,D. In adults low level expression is seen in many tissues with high levels of expression seen in a subset of sensory neurons in the head and in the spermatheca and somatic gonad.

### Table S3. GO term, network, and pathways analysis

| GO terms | | | | | |
|---|---|---|---|---|---|
| **term ID** | **term description** | **N obs** | **N bkg** | **strength** | **Q** |
| GO:0006635 | Fatty acid beta-oxidation | 8 | 29 | 0.95 | 0.0015 |
| GO:0000097 | Sulfur amino acid biosynthetic process | 7 | 26 | 0.94 | 0.0049 |
| GO:0072329 | Monocarboxylic acid catabolic process | 12 | 50 | 0.89 | 0.0000601 |
| GO:0009062 | Fatty acid catabolic process | 11 | 45 | 0.89 | 0.00013 |
| GO:0009066 | Aspartate family amino acid metabolic process | 7 | 29 | 0.89 | 0.0085 |
| GO:1901607 | Alpha-amino acid biosynthetic process | 11 | 59 | 0.78 | 0.001 |
| GO:0046395 | Carboxylic acid catabolic process | 18 | 99 | 0.77 | 0.00000389 |
| GO:0044242 | Cellular lipid catabolic process | 14 | 77 | 0.77 | 0.00011 |
| GO:0050830 | Defense response to gram-positive bacterium | 13 | 70 | 0.77 | 0.0002 |
| GO:0016042 | Lipid catabolic process | 19 | 113 | 0.73 | 0.00000429 |
| GO:0002376 | Immune system process | 35 | 230 | 0.69 | 1.46E-10 |
| GO:0045087 | Innate immune response | 34 | 224 | 0.69 | 3.05E-10 |
| GO:0006631 | Fatty acid metabolic process | 16 | 104 | 0.69 | 0.00012 |
| GO:0051707 | Response to other organism | 49 | 339 | 0.67 | 1.07E-13 |
| GO:0009617 | Response to bacterium | 23 | 158 | 0.67 | 0.0000019 |
| GO:0050829 | Defense response to gram-negative bacterium | 14 | 99 | 0.66 | 0.0011 |
| GO:0098542 | Defense response to other organism | 47 | 336 | 0.65 | 3.06E-13 |
| GO:0042742 | Defense response to bacterium | 22 | 157 | 0.65 | 0.00000604 |
| GO:0044282 | Small molecule catabolic process | 19 | 139 | 0.64 | 0.000066 |
| GO:0032787 | Monocarboxylic acid metabolic process | 20 | 151 | 0.63 | 0.000056 |
| GO:1901605 | Alpha-amino acid metabolic process | 16 | 119 | 0.63 | 0.00046 |
| GO:0046394 | Carboxylic acid biosynthetic process | 14 | 113 | 0.6 | 0.0035 |
| GO:0019752 | Carboxylic acid metabolic process | 39 | 339 | 0.57 | 9.21E-09 |
| GO:0006520 | Cellular amino acid metabolic process | 20 | 177 | 0.56 | 0.00035 |
| GO:0044283 | Small molecule biosynthetic process | 21 | 193 | 0.54 | 0.00035 |
| GO:0006082 | Organic acid metabolic process | 44 | 415 | 0.53 | 6.62E-09 |

| | | | | | |
|---|---|---|---|---|---|
| GO:0055114 | Oxidation-reduction process | 65 | 647 | 0.51 | 1.32E-12 |
| GO:0006790 | Sulfur compound metabolic process | 14 | 140 | 0.51 | 0.0241 |
| GO:0009605 | Response to external stimulus | 56 | 596 | 0.48 | 0.000000 001 |
| GO:0044281 | Small molecule metabolic process | 60 | 712 | 0.43 | 8.78E-09 |
| GO:0044255 | Cellular lipid metabolic process | 27 | 337 | 0.41 | 0.0022 |
| GO:0006629 | Lipid metabolic process | 34 | 432 | 0.4 | 0.00035 |
| GO:0006950 | Response to stress | 68 | 881 | 0.39 | 1.24E-08 |
| GO:0070887 | Cellular response to chemical stimulus | 31 | 404 | 0.39 | 0.0013 |
| GO:0044248 | Cellular catabolic process | 46 | 671 | 0.34 | 0.00024 |
| GO:1901575 | Organic substance catabolic process | 47 | 696 | 0.34 | 0.00026 |
| GO:0009056 | Catabolic process | 54 | 804 | 0.33 | 0.000066 |
| GO:0008152 | Metabolic process | 198 | 4808 | 0.12 | 0.0015 |
| GO:0031406 | Carboxylic acid binding | 7 | 33 | 0.83 | 2.26E-02 |
| GO:0005506 | Iron ion binding | 19 | 120 | 0.71 | 2.35E-05 |
| GO:0016614 | Oxidoreductase activity, acting on ch-oh group of donors | 10 | 64 | 0.7 | 0.0126 |
| GO:0004553 | Hydrolase activity, hydrolyzing o-glycosyl compounds | 9 | 65 | 0.65 | 0.0403 |
| GO:0016705 | Oxidoreductase activity, acting on paired donors, with incorporation or reduction of molecular oxygen | 17 | 130 | 0.62 | 0.00066 |
| GO:0046906 | Tetrapyrrole binding | 19 | 154 | 0.6 | 0.00045 |
| GO:0004497 | Monooxygenase activity | 12 | 100 | 0.58 | 0.0208 |
| GO:0020037 | Heme binding | 17 | 152 | 0.55 | 0.0033 |
| GO:0016491 | Oxidoreductase activity | 54 | 520 | 0.52 | 1.34E-10 |
| GO:0008194 | UDP-glycosyltransferase activity | 15 | 153 | 0.5 | 0.0226 |
| GO:0003824 | Catalytic activity | 215 | 3986 | 0.24 | 1.50E-13 |
| GO:0016787 | Hydrolase activity | 87 | 1617 | 0.24 | 0.0004 |
| GO:0043169 | Cation binding | 82 | 1670 | 0.2 | 0.0092 |
| GO:0046872 | Metal ion binding | 80 | 1655 | 0.19 | 0.0154 |
| GO:0043167 | Ion binding | 129 | 2786 | 0.17 | 0.001 |
| GO:0005764 | Lysosome | 14 | 98 | 0.66 | 0.007 |

| | | | | | |
|---|---|---|---|---|---|
| GO:0005737 | Cytoplasm | 150 | 3594 | 0.13 | 0.0374 |
| Local Network Cluster (String) | | | | | |
| CL:22702 | Mixed, incl. thiamine transporter 1, and abc transporter transmembrane region 2 | 5 | 5 | 1.51 | 0.00082 |
| CL:6159 | Hsp20/alpha crystallin family, and protein folding chaperone | 5 | 6 | 1.43 | 0.0013 |
| CL:15620 | Drug catabolic process | 4 | 5 | 1.41 | 0.0096 |
| CL:30955 | Aminoacylase activity, and cysteine-rich hydrophobic domain-containing protein 1/2 | 4 | 5 | 1.41 | 0.0096 |
| CL:22402 | Mixed, incl. sodium:potassium-exchanging atpase activity, and response to gamma radiation | 4 | 6 | 1.33 | 0.0133 |
| CL:23937 | Mixed, incl. methyltransferase domain, and neuropeptide | 4 | 6 | 1.33 | 0.0133 |
| CL:8140 | Cobalamin metabolic process, and methylmalonyl-coa mutase | 4 | 6 | 1.33 | 0.0133 |
| CL:8314 | Mixed, incl. fumarylacetoacetase-like, c-terminal, and negative regulation of map kinase activity | 4 | 6 | 1.33 | 0.0133 |
| CL:22685 | Mostly uncharacterized, incl. thiamine transporter 1, and pre-set motif | 7 | 14 | 1.2 | 0.00034 |
| CL:22388 | Mixed, incl. cub-like domain, and sodium:potassium-exchanging atpase activity | 6 | 13 | 1.17 | 0.0019 |
| CL:7823 | enoyl-CoA hydratase activity, and Acyl-CoA dehydrogenase, conserved site | 7 | 16 | 1.15 | 0.00064 |
| CL:30953 | Mixed, incl. aminoacylase activity, and phenazine biosynthesis-like protein | 5 | 13 | 1.09 | 0.0133 |
| CL:8330 | Mixed, incl. lactate metabolic process, and aldo/keto reductase family | 6 | 16 | 1.08 | 0.0042 |
| CL:23919 | Mostly uncharacterized, incl. anion transporter sulp-4/5, and c-terminus of aa_permease | 5 | 15 | 1.03 | 0.0188 |
| CL:22387 | Mixed, incl. cub-like domain, and nematode fatty acid retinoid binding protein (gp-far-1) | 8 | 25 | 1.01 | 0.00077 |
| CL:23918 | Mostly uncharacterized, incl. defense response to fungus, and anion transporter sulp-4/5 | 7 | 25 | 0.95 | 0.0043 |
| CL:34856 | Uncharacterised protein family UPF0376, and Putative pyroglutamyl peptidase 1 | 5 | 18 | 0.95 | 0.0363 |
| CL:22386 | Mixed, incl. cub-like domain, and transmembrane glycoprotein | 11 | 40 | 0.94 | 0.0000837 |
| CL:20327 | Mostly uncharacterized, incl. blood group rhesus c/e/d polypeptide, and glycolipid transfer protein domain | 6 | 22 | 0.94 | 0.0133 |
| CL:23917 | Mixed, incl. metapathway udp-glucuronosyltransferases, and defense response to fungus | 8 | 30 | 0.93 | 0.0019 |

| | | | | | |
|---|---|---|---|---|---|
| CL:22174 | Mixed, incl. innate immune response, and membrane raft | 32 | 146 | 0.85 | 1.31E-12 |
| CL:7821 | Fatty acid beta-oxidation, and acyl-coa dehydrogenase, n-terminal domain | 10 | 45 | 0.85 | 0.00079 |
| CL:7819 | Fatty acid beta-oxidation, and acyl-coa dehydrogenase/oxidase c-terminal | 12 | 57 | 0.83 | 0.00021 |
| CL:22181 | Mostly uncharacterized, incl. md domain, and membrane raft | 12 | 58 | 0.82 | 0.00022 |
| CL:22176 | Mostly uncharacterized, incl. membrane raft, and papain family cysteine protease | 21 | 105 | 0.81 | 0.000000142 |
| CL:22177 | Mixed, incl. membrane raft, and papain family cysteine protease | 17 | 85 | 0.81 | 0.0000063 |
| CL:7817 | Valine, leucine and isoleucine degradation, and Acyl-CoA dehydrogenase-like, C-terminal | 15 | 74 | 0.81 | 0.0000263 |
| CL:22180 | Mostly uncharacterized, incl. membrane raft, and md domain | 13 | 67 | 0.79 | 0.00018 |
| CL:22184 | Mixed, incl. md domain, and membrane raft | 9 | 47 | 0.79 | 0.0046 |
| CL:21371 | Mixed, incl. up-regulated in daf-2, and protein of unknown function (duf684) | 6 | 31 | 0.79 | 0.0495 |
| CL:22173 | Mixed, incl. innate immune response, and membrane raft | 34 | 179 | 0.78 | 3.24E-12 |
| CL:7814 | Valine, leucine and isoleucine degradation, and Fatty acid metabolism | 17 | 91 | 0.78 | 0.0000131 |
| CL:22179 | Mixed, incl. membrane raft, and cysteine peptidase, histidine active site | 14 | 74 | 0.78 | 0.000097 |
| CL:15602 | Cytochrome P450, E-class, group I | 8 | 47 | 0.74 | 0.0169 |
| CL:23845 | Mixed, incl. defense response to fungus, and defense response to gram-negative bacterium | 11 | 66 | 0.73 | 0.0024 |
| CL:15604 | Drug catabolic process, and cytochrome p450, e-class, group i | 7 | 42 | 0.73 | 0.0427 |
| CL:20259 | Mixed, incl. defense response to gram-positive bacterium, and blood group rhesus c/e/d polypeptide | 9 | 56 | 0.71 | 0.0129 |
| CL:8778 | Mixed, incl. glutathione metabolism, and cellular oxidant detoxification | 16 | 107 | 0.68 | 0.00021 |
| CL:7800 | Mixed, incl. fatty acid metabolism, and valine, leucine and isoleucine degradation | 19 | 130 | 0.67 | 0.0000462 |
| CL:34801 | Mixed, incl. uncharacterised protein family upf0376, and paw domain superfamily | 8 | 56 | 0.66 | 0.0431 |
| CL:8782 | Glutathione metabolism, and cellular oxidant detoxification | 11 | 80 | 0.64 | 0.0096 |
| CL:34794 | Mixed, incl. uncharacterised protein family upf0376, and glycoprotein catabolic process | 11 | 83 | 0.63 | 0.0118 |

| | | | | | |
|---|---|---|---|---|---|
| CL:34793 | Mixed, incl. domain of unknown function (duf19), and glycoprotein catabolic process | 13 | 119 | 0.54 | 0.0145 |
| CL:15592 | Mixed, incl. monooxygenase activity, and nuclear hormone receptor, ligand-binding domain | 21 | 199 | 0.53 | 0.00064 |
| CL:15595 | Mixed, incl. monooxygenase activity, and steroid metabolic process | 20 | 193 | 0.52 | 0.0011 |
| CL:7515 | Mixed, incl. carbon metabolism, and arginine and proline metabolism | 14 | 143 | 0.5 | 0.0213 |
| CL:15596 | Mixed, incl. monooxygenase activity, and steroid metabolic process | 16 | 173 | 0.47 | 0.0161 |
| KEGG Pathways | | | | | |
| cel00670 | One carbon pool by folate | 4 | 12 | 1.03 | 0.0124 |
| cel00280 | Valine, leucine and isoleucine degradation | 13 | 52 | 0.9 | 0.00000403 |
| cel00640 | Propanoate metabolism | 8 | 33 | 0.89 | 0.00073 |
| cel00220 | Arginine biosynthesis | 4 | 17 | 0.88 | 0.0289 |
| cel00062 | Fatty acid elongation | 6 | 27 | 0.85 | 0.0054 |
| cel00410 | beta-Alanine metabolism | 6 | 27 | 0.85 | 0.0054 |
| cel00071 | Fatty acid degradation | 10 | 51 | 0.8 | 0.00059 |
| cel00630 | Glyoxylate and dicarboxylate metabolism | 7 | 36 | 0.79 | 0.0046 |
| cel00260 | Glycine, serine and threonine metabolism | 5 | 27 | 0.77 | 0.0254 |
| cel00620 | Pyruvate metabolism | 5 | 29 | 0.74 | 0.0289 |
| cel01212 | Fatty acid metabolism | 11 | 67 | 0.72 | 0.00073 |
| cel01230 | Biosynthesis of amino acids | 11 | 71 | 0.7 | 0.00078 |
| cel00010 | Glycolysis / Gluconeogenesis | 6 | 40 | 0.68 | 0.0254 |
| cel00980 | Metabolism of xenobiotics by cytochrome P450 | 6 | 46 | 0.62 | 0.0362 |
| cel04142 | Lysosome | 12 | 102 | 0.58 | 0.0033 |
| cel01200 | Carbon metabolism | 12 | 111 | 0.54 | 0.0053 |
| cel01100 | Metabolic pathways | 69 | 916 | 0.38 | 9.86E-09 |
| **WikiPathways** | | | | | |
| WP148 | Fatty acid beta-oxidation 2 | 3 | 4 | 1.38 | 0.0229 |
| WP499 | Fatty acid beta-oxidation 3 | 3 | 7 | 1.14 | 0.026 |
| WP126 | Fatty acid beta-oxidation 1 | 5 | 20 | 0.9 | 0.0229 |
| WP38 | Fatty acid biosynthesis | 4 | 16 | 0.9 | 0.026 |
| WP209 | Fatty acid beta-oxidation meta-pathway | 5 | 22 | 0.86 | 0.0229 |

| | | | | | |
|---|---|---|---|---|---|
| WP1431 | Metapathway UDP-glucuronosyltransferases | 13 | 66 | 0.8 | 0.0000382 |
| WP1451 | Valine, leucine and isoleucine degradation | 6 | 32 | 0.78 | 0.0229 |
| **Monarch WBpheno** | | | | | |
| WBPhenotype:0001655 | Cadmium hypersensitive | 15 | 80 | 0.78 | 0.00014 |
| WBPhenotype:0001653 | Cadmium response variant | 16 | 91 | 0.75 | 0.00014 |
| WBPhenotype:0000591 | Metal response variant | 22 | 145 | 0.69 | 0.0000147 |
| WBPhenotype:0000012 | Dauer constitutive | 16 | 123 | 0.62 | 0.0025 |
| WBPhenotype:0000136 | mRNA levels increased | 15 | 149 | 0.51 | 0.048 |
| WBPhenotype:0001918 | Chemical hypersensitive | 29 | 369 | 0.4 | 0.0053 |
| **UniProt Keywords** | | | | | |
| KW-0442 | Lipid degradation | 8 | 40 | 0.81 | 0.0051 |
| KW-0503 | Monooxygenase | 12 | 91 | 0.63 | 0.0051 |
| KW-0349 | Heme | 14 | 122 | 0.57 | 0.0051 |
| KW-0408 | Iron | 21 | 194 | 0.54 | 0.00033 |
| KW-0443 | Lipid metabolism | 15 | 153 | 0.5 | 0.0091 |
| KW-0560 | Oxidoreductase | 30 | 312 | 0.49 | 0.0000614 |
| KW-0378 | Hydrolase | 48 | 805 | 0.28 | 0.0029 |
| KW-0732 | Signal | 181 | 4029 | 0.16 | 0.0000614 |
| **Pfam** | | | | | |
| PF02408 | CUB-like domain | 10 | 50 | 0.81 | 0.0047 |
| PF00067 | Cytochrome P450 | 14 | 76 | 0.77 | 0.001 |
| PF00201 | UDP-glucoronosyl and UDP-glucosyl transferase | 13 | 75 | 0.74 | 0.0024 |
| PF04101 | Glycosyltransferase family 28 C-terminal domain | 12 | 71 | 0.73 | 0.0047 |
| PF01579 | Domain of unknown function (DUF19) | 13 | 84 | 0.7 | 0.0047 |
| PF13561 | Enoyl-(Acyl carrier protein) reductase | 11 | 81 | 0.64 | 0.0345 |
| PF00106 | Short chain dehydrogenase | 11 | 82 | 0.63 | 0.0345 |
| **InterPro** | | | | | |

| | | | | | |
|---|---|---|---|---|---|
| IPR036264 | Bacterial exopeptidase dimerisation domain | 4 | 5 | 1.41 | 0.0273 |
| IPR002053 | Glycoside hydrolase, family 25 | 5 | 9 | 1.25 | 0.0156 |
| IPR020904 | Short-chain dehydrogenase/reductase, conserved site | 8 | 38 | 0.83 | 0.0204 |
| IPR003366 | CUB-like domain | 10 | 50 | 0.81 | 0.0058 |
| IPR001128 | Cytochrome P450 | 14 | 76 | 0.77 | 0.0009 |
| IPR036396 | Cytochrome P450 superfamily | 14 | 76 | 0.77 | 0.0009 |
| IPR002213 | UDP-glucuronosyl/UDP-glucosyltransferase | 13 | 71 | 0.77 | 0.0012 |
| IPR002401 | Cytochrome P450, E-class, group I | 13 | 73 | 0.76 | 0.0013 |
| IPR017972 | Cytochrome P450, conserved site | 12 | 71 | 0.73 | 0.0043 |
| IPR002542 | Domain of unknown function DUF19 | 11 | 72 | 0.69 | 0.0156 |
| IPR029058 | Alpha/Beta hydrolase fold | 24 | 187 | 0.61 | 0.0001 |
| IPR036291 | NAD(P)-binding domain superfamily | 17 | 153 | 0.55 | 0.0074 |
